# USP5 enhances SGTA mediated protein quality control

**DOI:** 10.1101/2021.09.13.460017

**Authors:** Jake Hill, Yvonne Nyathi

## Abstract

Mislocalised membrane proteins (MLPs) present a risk to the cell due to exposed hydrophobic amino acids which cause MLPs to aggregate. Previous studies identified SGTA as a key component of the machinery that regulates the quality control of MLPs. Overexpression of SGTA promotes deubiqutination of MLPs resulting in their accumulation in cytosolic inclusions, suggesting SGTA acts in collaboration with deubiquitinating enzymes (DUBs) to exert these effects. However, the DUBs that play a role in this process have not been identified. In this study we have identified the ubiquitin specific peptidase 5 (USP5) as a DUB important in regulating the quality control of MLPs. We show that USP5 is in complex with SGTA, and this association is increased in the presence of an MLP. Overexpression of SGTA results in an increase in steady-state levels of MLPs suggesting a delay in proteasomal degradation of substrates. However, our results show that this effect is strongly dependent on the presence of USP5. We find that in the absence of USP5, the ability of SGTA to increase the steady state levels of MLPs is compromised. Moreover, knockdown of USP5 results in a reduction in the steady state levels of MLPs, while overexpression of USP5 increases the steady state levels. Our findings suggest that the interaction of SGTA with USP5 enables specific MLPs to escape proteasomal degradation allowing selective modulation of MLP quality control. These findings progress our understanding of aggregate formation, a hallmark in a range of neurodegenerative diseases and type II diabetes, as well as physiological processes of aggregate clearance.

## Introduction

Newly synthesised membrane proteins are triaged between biosynthetic pathways targeting them to subcellular compartments such as the mitochondria and the endoplasmic reticulum (ER) and degradative pathways operating in the cytosol [1]. Timely degradation of membrane proteins that mislocalise to the cytosol is necessary to prevent their accumulation and aggregation which is linked to a range of pathologies [2, 3]. Therefore, a variety of intricate protein quality control mechanisms are dedicated to eliminating these mislocalised membrane proteins (MLPs) and maintaining protein homeostasis in mammalian cells [4]. Another layer of protein quality control for membrane proteins occurs at the ER membrane. Misfolded membrane proteins that fail to integrate into the ER membrane are targeted for ubiquitination and subsequent degradation via the endoplasmic-reticulum-associated protein degradation pathway (ERAD) [5, 6]). These quality control mechanisms collectively act to ensure that mislocalised or misfolded membrane proteins are degraded at any stage during their targeting and integration into the ER membrane [7].

The fate of MLPs that expose hydrophobic amino acids in the cytosol is decided via a delicate collaboration between factors that promote ubiquitination and those that promote deubiquitination of the MLPs [8, 9]. Ubiquitination occurs via the action of an array of enzymes; an E1 ubiquitin activating enzyme, an E2 ubiquitin-conjugating enzyme and an E3 ubiquitin ligase, acting in sequence to polyubiquitinate the substrate for subsequent proteasomal degradation [10]. Ubiquitination is a highly dynamic and reversible process due to the action of deubiquitinating enzymes (DUBs) which act as ubiquitin proteases, thereby antagonising the action of the E3 ligases and preventing the proteasomal degradation of substrates [11], [12].

It has long been known that DUBs are important for generating a pool of free ubiquitin in the cell [13],[14]. However, recent evidence suggests a role for DUBs in the regulation of many cellular processes by fine-tuning or modulating substrate ubiquitination and degradation [14]. A typical example is the DUB Ataxin-3, which acts to limit the length of ubiquitin chains built on substrates by CHIP (C-terminus of Hsc70 Interacting Protein) a mammalian E3 ligase involved in cytosolic protein quality control [15]. In addition, some cytosolic localised DUBs are important in kinetic proofreading and discrimination of membrane proteins that should be degraded by the proteasome [16]. While DUBs localised on the proteasome, such as UCHL5 and USP14 can delay the degradation of proteasomal substrates [17]. Moreover, DUBs are now known to have an important role in ubiquitin chain editing by promoting either K48 ubiquitin conjugates or K63 linked ubiquitin chains on substrates to regulate degradation by the proteasome or via autophagy, respectively [18, 19].

Recent studies suggest a role for USP13 in ERAD [20, 21], and USP20/33 in deubiquitinating tail anchored membrane proteins prior to their insertion into the ER membrane [22]. Although such details about the role of specific DUBs in membrane protein quality control is beginning to emerge, this knowledge in still in its infancy. This is in stark contrast to the tremendous progress that has been made towards our understanding of the E3 ligases that play a role in membrane protein quality control. In yeast the cytosolic ubiquitin ligases Ubr1p [23], San1P [24] and the ER resident Doa10p and Hrd1p ubiquitinate substrates for proteasomal degradation [25, 26]. The mammalian orthologues of Hrd1p (HRD1) [27], Hrd1p homologue gp78 [28] and Doa10p (MARCH6) [29] are known to promote ERAD. Moreover, the ligases MARCH6 and TRC8 were also shown to promote the ubiquitination of tail anchored proteins at the cytosolic face of the ER [30], while RNF126 was shown to promote ubiquitination of MLPs [31].

Several studies have also demonstrated a role for the ubiquitin proteasome system in the degradation of MLPs [4, 9, 31-35]. Among the factors known to be involved in regulating this degradation is the BCL2-associated athanogene 6 (BAG6) [1, 36, 37], the cochaperone SGTA (small glutamine-rich tetratricopeptide repeat protein alpha) [1, 9, 32] and a host of their diverse interacting partners. SGTA, is a homodimer, with an N-terminal domain which interacts with UBL4A (the ubiquitin-like protein 4A), and BAG6 [38, 39], a central TPR domain which interacts with Hsp70 and Hsp90 chaperones [40, 41], and a C-terminal domain capable of binding hydrophobic substrates [42].

A BAG6/SGTA quality control cycle dictating the fate of MLPs is proposed to be operating at the proteasome [43]. In this cycle, BAG6 recruits RNF126, an E3 ubiquitin ligase which catalyses the selective ubiquitination of MLPs for subsequent degradation by the proteasome [31, 44]. On the other hand, SGTA promotes deubiquitination thereby giving substrates a chance to escape proteasomal degradation [9, 32]. Therefore, in the context of the BAG6/SGTA cycle, it is plausible that an interaction of SGTA with deubiquitinating enzymes could provide SGTA-bound MLPs access to the enzyme(s) which mediate deubiquitination, thereby delaying the clearance of the MLPs and perhaps favour their targeting to the ER and subsequent membrane integration [43]. However, the deubiquitinating enzyme(s) that play a role in this process have not been defined to date. In this study we employed pull downs and mass spectrometry to identify DUBs that interact with SGTA. We show that USP5 is in complex with SGTA using co-immunoprecipitation, and that this interaction is necessary for the ability of SGTA to increase the steady state levels of MLPs. The implications of this study to our understanding of mislocalised membrane protein quality control is discussed.

## Materials and methods

All cell culture and standard reagents were purchased from Fisher Scientific (Loughborough, UK). Antibodies to opsin and V5 were previously described [32]. Commercially available antibodies against the following targets were purchased from the indicated suppliers, tubulin and GFP were purchased from Abcam (Cambridge, UK), USP9X (#A301-350A), USP5 (#A301-542A), USP14 (#A300-920A) and USP10 (#A300-900A) antibodies were purchased from Bethyl Laboratories Inc. (Montgomery, USA). Bortezomib was purchased from Selleckchem (Munich, Germany), Leupeptin and Pepstatin A were purchased from BIOMOL (Hamburg, Germany).

### Plasmids

Plasmids for the overexpression of SGTA-V5 and Pex19-V5 were previously described [43]. Ub-M–GFP (#11938) and Ub-R–GFP (#11939) plasmids described in [45] were purchased from Addgene. HA tagged USP5 (#22590) was purchased from Addgene and subcloned into pcDNA5 mammalian expression vector. The 3x NNP/AAA (NNP positions at 226-228, 239-241 and 255-257) variant of SGTA-V5 was previously described [46].

The pET28-TxA-SGTA_wild type plasmid for the expression and purification of WT SGTA was previously described [46]. This plasmid was generated by cloning WT SGTA into BamHI/XhoI restriction sites of a pET28c vector modified to encode an N-terminal thioredoxin A (TxA) fusion protein followed by a hexa-histidine tag. Plasmids for the expression and purification of SGTA_TPR double mutant (SGTA^K160E/R164E^) and SGTA_UBL double mutant (SGTA^D27R/E30R^) mutant were created by site directed mutagenesis using pET28-TxA-SGTA wild type plasmid as template and primers carrying the mutated codons. DNA sequencing was used to verify the mutants prior to use.

### Cell culture

The inducible stable HeLa Flp-In T-REx cell lines expressing OP91 or opsin degron (OpD) were previously described in [32] and [47], respectively. The HeLa cells were maintained in DMEM containing 10% fetal bovine serum and 2 mM L-glutamine at 37°C under 5% CO2. At alternate passages, DMEM was supplemented with 100 μg/ml hygromycin B and 4 μg/ml blasticidin S. Transfections of plasmid DNA were performed using Lipofectamine 2000 (Life Technologies, Carlsbad, CA, USA) in accordance with the manufacturer’s instructions. 6 hours post-transfection, OP91 or OpD expression was induced by treating the cells with DMEM containing 1 μg/ml tetracycline followed by growth for an additional 16-18 hours. Where indicated, 10 nM bortezomib or 100 µM leupeptin and 1 µg/ml pepstatin were added to cells 20 hours prior to analysis.

### Knockdown experiments

For siRNA experiments, 20 nM of siRNAi duplexes were transfected into cells using INTERFERin (Polyplus, Illkirch, France) according to the manufacturer’s instructions. 48 hours post-transfection the cells were transiently transfected with the appropriate DNA constructs (where necessary) and harvested after a further 24-hour incubation. Non-targeting control duplex [Cat# D-001810-01-20] and synthetic siRNA duplexes, USP5 siRNA1; [Cat#: D-006095-02-0002] and USP5 siRNA2; [Cat# D-006095-03-0002] targeting the USP5 gene [13], and the USP9x ON-TARGETplus siRNA [Cat# J-006099-06-0002] were purchased from Dharmacon (Chicago, IL, USA). SiRNA target sequences for SGTA and USP14 were 5’-ACAAGAAGCGCCUGGCCTATT-3’ [48] and 5’-AGAAATGCCTTGTATATCATT-3’ [49], respectively. SiRNA duplexes for these were obtained from Dharmacon (Chicago, IL, USA).

### Real-time RT-PCR

To quantify OP91 expression levels, total cellular RNAs were isolated using TRIzol™ Plus RNA Purification Kit (Invitrogen, Carlsbad, CA) and used for first-strand cDNA using the SensiFAST™ cDNA Synthesis (Bioline, Memphis, TN, USA). Quantitation of the OP91 transcript was done by quantitative PCR using the SensiFAST SYBR® Hi-ROX Kit (Bioline, Memphis, TN, USA) and the StepOnePlus™ Real-Time PCR detection system (Applied biosystems by Thermo Fisher Scientific). The expression of GAPDH was used as the internal control. For the primers 5 pmol of OP91 AGGGCCCAAACTTCTACGTG (forward) and AGCGTGAGGAAGTTGATGGG (reverse) and GAPDH, GAAGGTGAAGGTCGGAGTC (forward) and GAAGATGGTGATGGGATTTC (reverse) were used. PCR conditions were 50°C for 2 min, 95°C for 10 min, and then 40 cycles of 95°C for 15 s and 60°C for 1 min. Melt curve analysis was performed to verify the specificity of the reaction and cycle threshold values (Ct values) were used to calculate the -fold differences using the comparative Ct (ΔΔCt) method.

### Protein purification

Plasmids for protein expression were transfected into *E. coli* BL21 (DE3) strains and protein expression was induced by using 0.5 mM isopropyl-β-D-thiogalactopyranoside (IPTG) at OD600 ≈ 0.8, followed by overnight incubation at 22 °C. Cells were harvested and resuspended in lysis buffer (20 mM potassium phosphate, pH 8.0, 300 mM NaCl, 10 mM Imidazole, 50 mM Tris-HCl, 10% glycerol, supplemented with protease inhibitors -EDTA (Roche, Burgess Hill, UK), and 1 mM PMSF and lysed by sonication. Cell debris was removed by centrifugation, and the soluble fractions were purified using nickel affinity chromatography (Qiagen, Manchester, UK) and eluted with lysis buffer containing 300 mM imidazole, followed by dialysis into phosphate buffered saline pH 7.4. Gel filtration was carried out using a HiLoad 16/60 Superdex 200 column (GE Healthcare, Amersham, UK), previously equilibrated in buffer containing 10 mM potassium phosphate pH 6.0, 100 mM NaCl and 50 mM Tris-HCl. Proteins were concentrated using Vivaspin concentrators (Sartorius Stedin) and the sample purity and homogeneity was assessed by SDS-PAGE.

### Coupling of purified recombinant protein to resin

UltraLink Biosupport resin (Thermo Scientific, Pierce) were incubated with coupling buffer (0.6M Sodium Citrate, 0.1M MOPS, pH 7.5) and equimolar amounts of recombinant protein WT SGTA, SGTA mutants (SGTA^K160E/R164E^ and SGTAD^27R/E30R^) or Thioredoxin, overnight at 4°C, to allow covalent coupling via primary amines. 3M ethanolamine pH 9.0 was used to quench the reaction after coupling. Beads with coupled recombinant proteins were washed for 15 minutes with PBS and 1M NaCl in turn. Beads were further washed 3 times in PBS and re-suspended in PBS for storage at 4 °C.

### Pull-down assays

Beads coupled to recombinant WT SGTA, SGTA mutants (SGTA^K160E/R164E^ and SGTA^D27R/E30R^) or Thioredoxin were incubated with rabbit reticulocyte lysate or HeLa lysate overnight. Resins were washed three times in buffer containing 40mM HEPES-KOH, pH 7.5, 40mM potassium acetate, 5mM MgCl_2_ followed by incubation in Laemmli sample buffer at 70 °C for 10 minutes. Samples were analysed by SDS-PAGE and western blotting.

### Mass spectrometry analysis

Mass spectrometry was performed at the Sanford-Burnham Proteomics Facility, Sanford-Burnham Medical Research Institute, (La Jolla) using the method previously described [50]. Briefly, beads from pull down assays were resuspended in a buffer containing 8M urea and 50mM ammonium bicarbonate followed by reduction with Tris(2-carboxyethyl) phosphine, (TCEP), alkylation with iodoacetamide and digestion with mass spectrometry grade trypsin (Promega). Samples were washed with 50mM ammonium bicarbonate and formic acid was added, followed by desalting. Peptides were analysed by high-resolution and high-accuracy LC-MS/MS. The LC-MS/MS raw data of three technical replicates were combined and submitted to Sorcerer Enterprise v.3.5 release (Sage-N Research Inc.) with SEQUEST algorithm as the search program for peptide/protein identification. The LC-MS/MS raw data were also submitted to Integrated Proteomics Pipelines (IP2) Version IP2 1.01 (Integrated Proteomics Applications, Inc.) with ProLucid algorithm as the search program for peptide/protein identification. ProLucid and SEQUEST were both set up to also search the target-decoy ipi. Human.v3.73 protein database. The search results were viewed, sorted, filtered, and statically analysed by using comprehensive proteomics data analysis software, Peptide/Protein prophet v.4.02 (ISB) and DTA Select for proteins. The differential spectral count analysis was done by QTools, an open-source tool for automated differential peptide/protein spectral count analysis and Gene Ontology. The reviewed human UniProt database was used for identifying SGTA interacting partners.

### Preparation of HeLa lysate

HeLa M cells were plated in 10-cm dishes at 100% confluency and washed twice with PBS at 4°C. Following washes, cells were lysed in a buffer containing 20mM HEPES pH 7.4, 5mM MgCl_2_, 0.1M NaCl, 0.5% Triton x100, protease inhibitor and 1mM PMSF. Cells were harvested from the dish using a cell scrapper followed by incubation on ice for 10-15 minutes. The cell suspension was then centrifuged to remove cell debris at 13000 rpm at 4 °C for 20 min and the supernatant used for analysis.

### Western blotting

Western blot experiments were performed according to standard procedures previously described [32]. Briefly, cells were lysed directly into Laemmli sample buffer and subjected to SDS-PAGE followed by infrared immunoblotting as described previously. Fluorescent bands for specific proteins were quantified using Image Studio-Lite (Li-Cor Biosciences) employing data from at least three independent experiments expressed as the mean ± s.e.m. The statistical significance of the results was assessed by applying a student’s t-test using Prism 7 (GraphPad).

### Co-immunoprecipitation

Stable HeLa Flp-In T-REx cell lines expressing OP91 or OpD were grown as detailed. Half of the samples were induced while the other half remained as uninduced samples. Cell lysates were prepared by harvesting cells in solubilisation buffer containing 10 mM Tris-HCl pH 7.4, 150 mM NaCl, 0.5% n-dodecyl β-D-maltopyranoside (DDM) and 1 mM EDTA supplemented with protease inhibitor tablets (Roche) and 1 mM PMSF for 1 hour at 4°C. The lysates were centrifuged at 16,000 g for 20 minutes at 4°C to remove insoluble material and 10% of the supernatant was retained as the input sample. Approximately 500 μL of the clarified lysate was incubated with IgG control or SGTA antibodies (1–2 μg) for 12 h at 4 °C with constant rotation. Protein A Sepharose (GenScript Biotech, Piscataway USA) was used to precipitate complexes by adding a 50% slurry and incubating for further 2 h. Beads were then washed five times in lysis buffer followed by elution of immune complexes by resuspending the beads in 2× Laemmli sample buffer followed by immunoblotting with appropriate antibodies.

### Cycloheximide chase

Transfected HeLa Flp-In T-REx cells were treated with non-targeting siRNAi duplexes or siRNAi targeting USP5, USP9X, USP14, and SGTA as described above. Prior to harvesting transfected cells were treated with 100 µg/ml cycloheximide (Sigma, Aldrich) to inhibit protein synthesis. Cells were lysed directly into SDS-PAGE sample buffer at specific time-points followed by Western blotting analysis.

## Results

Previous studies suggest that SGTA regulates the fate of membrane proteins that mislocalise to the cytosol [9, 32]. Overexpression of exogenous SGTA led to an increase in the steady-state levels of MLPs. One such model MLP is OP91, an N-terminal fragment of opsin that is inefficiently targeted to the endoplasmic reticulum [32]. SGTA also promoted the deubiquitination of MLPs [9, 32]. Since, SGTA itself has no domains typical for DUBs, it was proposed that SGTA recruits one or more DUBs to exert these effects [9, 32, 43]. In this study we aimed to identify the DUBs that interact with SGTA in mammalian cells and dissect their role in MLP quality control.

### Identification of SGTA Interacting DUBs

To identify SGTA interacting DUBs, we took a proteomic based approach by isolating protein complexes that interact with SGTA using a pull-down assay. We made use of SGTA mutants that are defective in interacting with known SGTA binding partners to develop and validate a robust assay for the identification of specific SGTA interactors. The SGTA^K160E/R164E^ mutant which does not interact with Hsp70 [48] and the (SGTA^D27R/E30R)^ which does not interact with BAG6 and Ubl4A [20], were utilised in the pull-downs. WT SGTA, and the two SGTA mutants (SGTA^D27R/E30R^ and SGTA^K160E/R164E^) were purified as thioredoxin (TXN) fusion proteins and immobilised to UltraLink Biosupport beads. Protein complexes interacting with these immobilised proteins were isolated from rabbit reticulocyte lysate under native conditions, with recombinant TXN immobilised to beads as negative control. The pull-downs were validated by Western blotting analysis, probing with BAG6, Ubl4A and Hsp70 specific antibodies. As expected, WT SGTA pulled-down high levels of BAG6, Ubl4A and Hsp70, while the TXN control did not pull-down any significant amounts of these three proteins (Fig. 1A, cf. lanes 2 and 3). Consistent with previous publications [20], very low levels of BAG6 and Ubl4A were detected in the SGTA^D27R/E30R^ mutant and Hsp70 levels were low in the SGTA^K160E/R164E^ mutant (Fig. 1A, cf. lanes 2, 3 and 4) [48].

**Fig 1.**
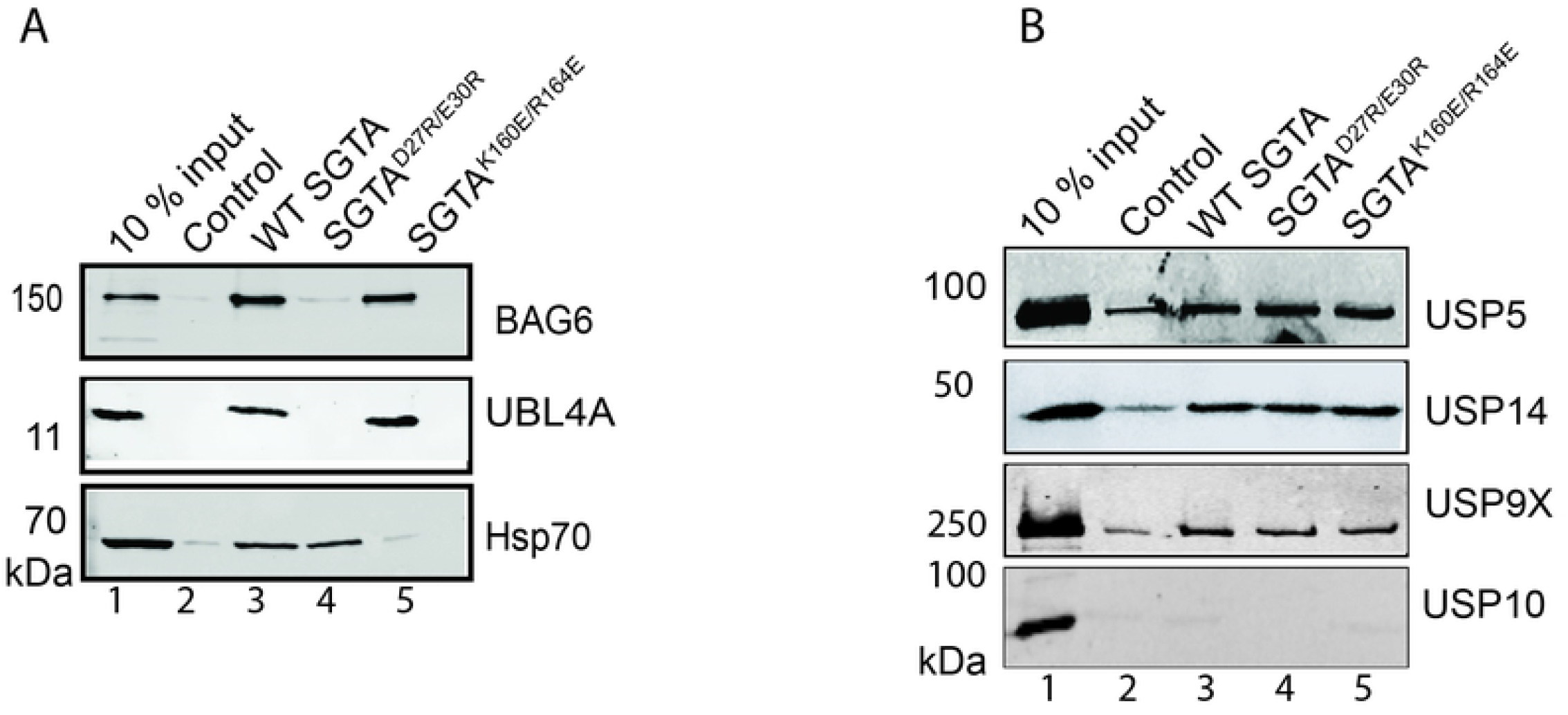
Identification of SGTA Interacting DUBs. **A)** Beads coupled to recombinant Thioredoxin-SGTA fusion proteins as indicated or Thioredoxin only (Control) were incubated with rabbit reticulocyte lysate for 16 hours at 4°C. The beads were washed as detailed in Materials and Methods. Twice the bead volume of SDS sample buffer was added, and samples were processed for SDS PAGE and Western blotting and probed with antibodies against Bag6, UBL4A and Hsp70 proteins using a fluorescence-based Odyssey ® Fc Imaging System (LICOR). **B)** Beads described in A, were incubated with HeLa lysate for 16 hours, washed and processed for Western blotting as detailed in A. Blots were probed with antibodies against USP5, USP14, USP9X and USP10 proteins using fluorescence- based detection on an Odyssey ® Fc Imaging System (LICOR).

Having validated that the pull-down approach was working as expected, pull-downs were performed as detailed above and the samples were analysed by mass spectrometry followed by interrogating the reviewed human UniProt database to identify SGTA interactors. Consistent with the Western blotting results (Fig. 1 A), beads containing WT SGTA were able to pull-down high levels of Bag6, Ubl4A and Hsp70 (Table 1) as judged from the spectral counts. The SGTA^D27R/E30R^ mutant had very low spectral counts for Bag6 and Ubl4A, while the SGTA^K160E/R164E^ mutant had very low levels of some isoforms of Hsp70, notably HspA2 known to specifically interact with SGTA [51]. Based on these results we gained confidence that this proteomic based approach was reliable and likely to reflect *bona fide* and novel SGTA interacting partners.

**Table 1.** SGTA interacting partners identified by mass spectrometry. Pull-down experiments were performed as described in Materials and Methods. The beads were then analysed for bound proteins by mass spectrometry as detailed in the materials and methods followed by interrogating the reviewed human UniProt database for SGTA interacting partners. Identified proteins known to interact with SGTA are indicated together with the spectral counts. SGTA peptides detected reflect comparative efficiency for the coupling of SGTA or the mutants to beads.

| Spectral counts |  |  |  |  |
| --- | --- | --- | --- | --- |
| SGTA interacting proteins | Thrx | SGTA | SGTA<sup>D27R/E30R</sup> | SGTA<sup>K160E/R164E</sup> |
| SGTA | 28.8794 | 1813.40948 | 1799.50341 | 1561.96154 |
| BAG6 | 70.6523 | 188.85411 | 26.81825 | 192.46266 |
| HSPA2 | 96.34687 | 37.44345 | 34.27598 | 0 |
| UBL4A | 10.9873 | 15.07829 | 2.7107 | 17.22824 |
| Number of SGTA peptides detected | 28 | 1813 | 1753 | 1799 |

We then looked at all the DUBs that had been identified by mass spectrometry (Table 2). Based on spectra counts, USP5, USP9X, USP14, and USP7 were identified as candidates for follow-up studies (Table 2). Since our goal was to focus on the DUBs that play a role in the quality control of membrane proteins mislocalised to the cytosol, we looked in published literature for the subcellular localisation of the DUBs that we had identified. Based on this criteria USP7 was not included in further analysis as it is reported to localise mainly to the nucleus, while USP5, USP9X and USP14 localise to both the nucleus and cytosol [52].

**Table 2.** Putative SGTA interacting DUBs identified by mass spectrometry. Pull-down experiments were performed as described in Materials and Methods. The beads were then analysed for bound proteins by mass spectrometry as detailed in the materials and methods. All DUBs identified from the mass spectrometry are shown together with the spectral counts. The DUBs highlighted were prioritised for further studies based on high spectral counts and low background in the negative control.

| Name of enzyme | Spectral |  |  |  |
| --- | --- | --- | --- | --- |
|  | Thrx | SGTA WT | SGTA<sup>D27R/E30R</sup> | SGTA<sup>K160E/R164E</sup> |
| Ubiquitin carboxyl-terminal hydrolase 10 (USP10) | 0 | 0 | 0 | 0.36738 |
| Ubiquitin carboxyl-terminal hydrolase 19 (USP19) | 0 | 0 | 0.68318 | 0 |
| **Ubiquitin carboxyl-terminal hydrolase 5 (USP5)** | **2.11094** | **9.27228** | **10.51442** | **8.43043** |
| **Ubiquitin carboxyl-terminal hydrolase 14 (USP14)** | **0** | **8.48274** | **14.24989** | **7.69728** |
| Ubiquitin carboxyl-terminal hydrolase 15 (USP15) | 6.86995 | 7.72786 | 7.81252 | 6.96413 |
| **Ubiquitin carboxyl-terminal hydrolase 7 (USP7)** | **0** | **6.18345** | **5.75859** | **3.66414** |
| Ubiquitin carboxyl-terminal hydrolase 47 (USP47) | 0 | 1.5502 | 1.35755 | 0 |
| Ubiquitin carboxyl-terminal hydrolase 32 (USP32) | 0 | 1.53864 | 0 | 0.36577 |
| Ubiquitin carboxyl-terminal hydrolase 4 (USP4) | 0 | 1.1612 | 1.02477 | 0 |
| Ubiquitin carboxyl-terminal hydrolase 24 (USP24) | 0 | 0.77221 | 1.69914 | 0 |
| **Probable ubiquitin carboxyl-terminal hydrolase FAF-X (USP9X)** | **0** | **5.40546** | **6.45057** | **4.39891** |
| Ubiquitin carboxyl-terminal hydrolase isozyme L5 (UCHL5) | 0 | 1.93919 | 1.69474 | 0.36738 |
| Ubiquitin thioesterase (OTUB1) | 0 | 1.93341 | 1.35755 | 0 |

Since the initial pull-downs which identified, these DUBs were performed using rabbit reticulocyte lysate we validated these results by repeating the pulldowns using lysate from HeLa cells. Pull-downs were performed as described above followed by Western blotting for the identified DUBs and USP10 which had very low spectral counts from the mass spectrometry results (Table 2). Consistent with the mass spectrometry data, WT SGTA and both mutants were able to pull down comparable amounts of USP5, USP9X, and USP14 when compared to TXN control, while USP10 was barely detectable, consistent with the very low spectral counts obtained for this DUB (Fig. 1B, cf. lanes 2, 3, 4 and 5). From these results we decided to further investigate the role of USP5, USP9X, and USP14 in SGTA mediated quality control of MLPs.

### Knocking down USP5 reduces the steady state of a model MLP

To investigate whether the identified DUBs play a role in MLP quality control, we reduced the level of endogenous DUBs using an siRNA approach and studied the effect on steady-state levels of the model MLP, OP91. We reasoned that if a DUB is responsible for the deubiquitination and stabilisation of OP91, its knock down would enhance MLP degradation through favouring the ubiquitinated form which is subject to proteasomal clearance. Knocking down USP9X did not reduce the steady state levels of OP91 when compared to the non-targeting siRNAi (Fig. 2A, lane 4), instead it led to a slight increase in the steady state levels of OP91 when signals were quantified and normalised to tubulin loading control (Figure 2B). Knocking down USP5 on the contrary led to a marked decrease in the steady state levels of non-glycosylated form of OP9, indicating a role on MLPs (Fig. 2A, cf. lane 1 and 2). The significant changes in the OP91 signal were equivalent to ∼50% reduction when compared to the non -targeting control (Figure 2B). Knockdown of USP14 also led to a decrease in the steady state levels of OP91, although the effect was less marked than that of USP5 (Fig. 2A, cf. lane 1, 2 and 3), possibly because the knockdown of USP14 was less efficient. In these studies, knocking down any of these DUBs had no effect on SGTA levels (Fig 2A), suggesting these results are directly related to changes in levels of the DUBs. The effect of these DUBs was specific for the non-glycosylated form of OP91, indicating that they act on MLPs.

**Fig 2.**
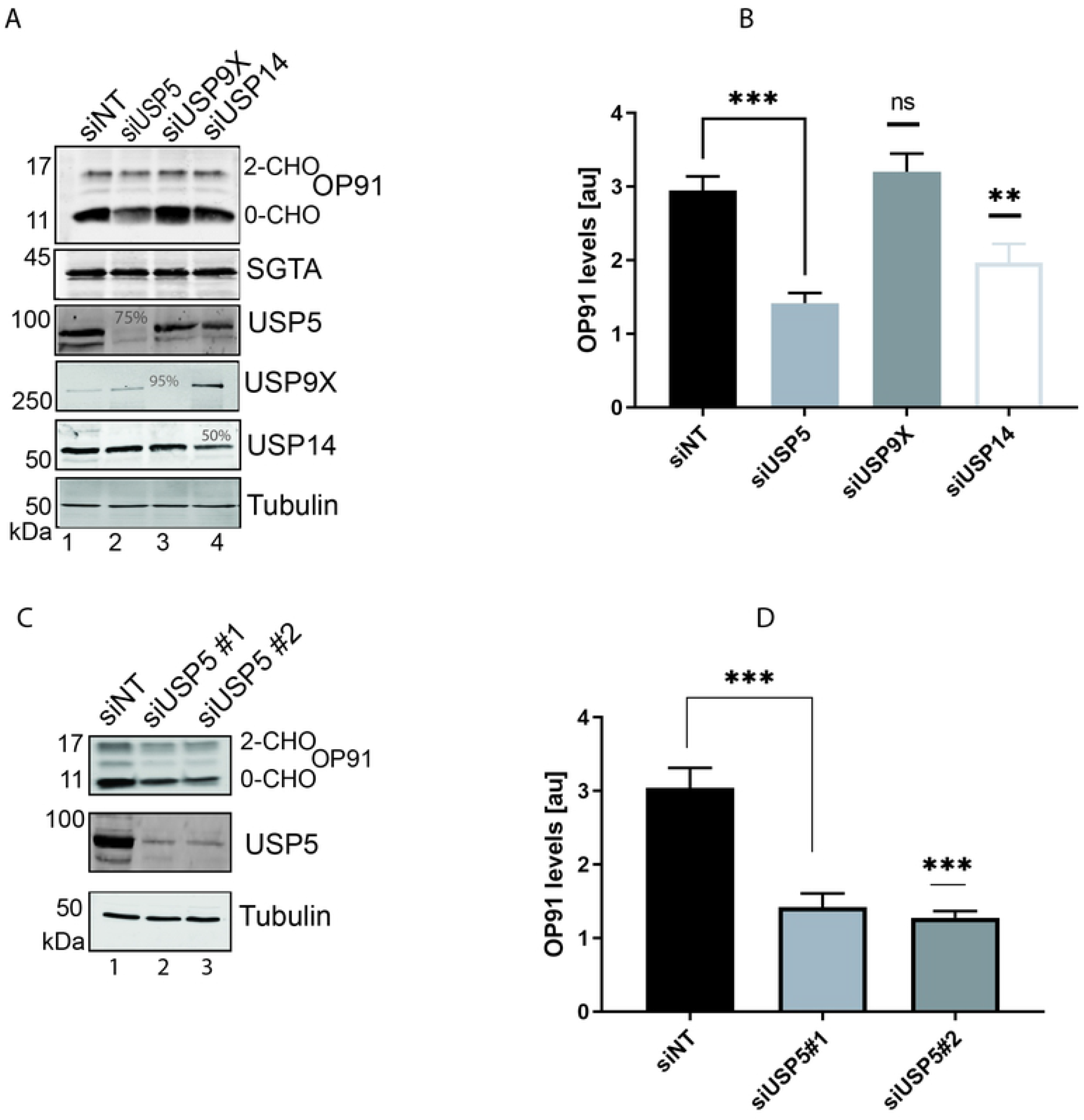
Knocking down USP5 reduces the steady state of a model MLP. **A)** HeLa TRex Flp-In cells stably expressing OP91 after induction, were seeded at 50 % confluence and transfected with 1 nM of siRNAi targeting the indicated DUBs. OP91 expression was induced 48 h later and cells grown for 20 hrs post induction. Total cell lysates were prepared, and products analysed by Western blotting with antibodies against OP91, SGTA, USP5, USP9X, USP14 and tubulin (loading control) using fluorescence-based detection (LICOR). The % knockdown efficiency is indicated for each DUB in lane 2, 3 and 4. **B**) OP91 signals were quantified using Odyssey 2.1 software and normalised to the tubulin loading control, values show standard errors for n = 3. Pairwise comparisons relative to the NT control, were performed using GraphPad Prism. Student t-test: P > 0.05 = ns, P ≤ 0.05= *, P ≤ 0.01= **, P ≤ 0.001= *** **C)** HeLa TRex Flp-In cells stably expressing OP91 were transfected with non-targeting siRNAi or two independent siRNAi duplexes targeting USP5. Cells were treated as described in A, followed by blotting for OP91, USP5 and tubulin. **D)** OP91 signals were quantified using Odyssey ® Fc Imaging system and normalised to the tubulin loading control, values show standard errors for n = 3. Pairwise comparisons relative to the NT control, were performed using GraphPad Prism. Student t-test: P > 0.05 = ns, P ≤ 0.05= *, P ≤ 0.01= **, P ≤ 0.001= *** and P ≤ 0.0001=****.

Mammalian USP14 and the yeast homologue, Ubp6 both associate with the proteasome [53], [54] and catalyse cleavage of ubiquitin subunits from substrates thereby delaying their proteasomal degradation [55]. OP91 was previously shown to be degraded by the proteasome [9], [32]. Hence, the destabilisation of OP91 upon knocking down USP14 is consistent with the effect of USP14 on proteasomal degradation of substrates. The destabilisation of MLPs upon USP5 knockdown is a novel finding that has not been previously reported. Hence, we focussed our attention on the role of USP5 in MLP quality control.

To validate the effects of USP5 on reducing the steady state levels of OP91, we made use of another siRNA oligonucleotide to independently target USP5. This siRNA oligonucleotide also led to a marked decrease in the level of OP91 (Fig. 2C). The effect of reducing OP91 levels was reproducible as both siRNA oligonucleotides targeting USP5 led to ∼ 50% reduction in OP91 levels (Fig. 2D).

### Loss of USP5 does not affect other substrates

Next, we wanted to confirm that the destabilisation effect observed after knocking down USP5 was specific for MLPs and not due to changes in ubiquitin homeostasis. To this effect, we made use of ubiquitin–methionine–GFP (Ub-M-GFP) which upon cleavage of ubiquitin does not have a degradation signal and is therefore stable [45]. USP9X and USP14 were included in this experiment for comparison. Knocking down any of the DUBs did not affect the steady state levels of Ub-M-GFP (Fig. 3A and 3B). We also made use of ubiquitin– arginine–GFP (Ub-R-GFP) [45]), a proteasomal N-end rule substrate whose degradation was previously shown to be independent of SGTA [43]. Knocking down all the DUBs led to comparable steady-state levels of Ub-R-GFP (Fig. 3A and 3B), suggesting the effects observed with USP5 are specific for OP91.

**Fig 3.**
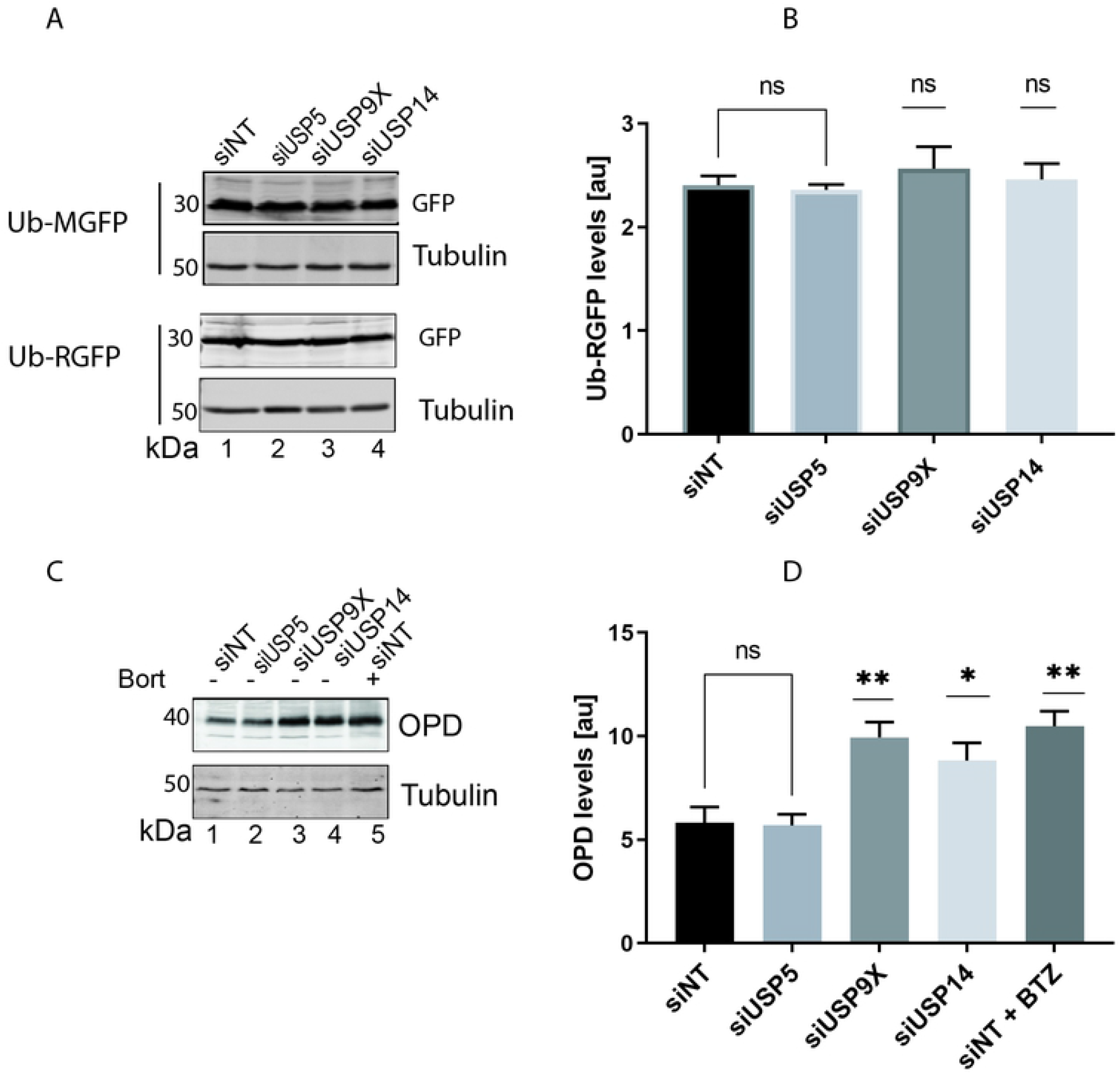
Loss of USP5 does not affect other substrates. **A)** HeLa M cells were seeded at 50 % confluence and transfected with 1 nM of siRNAi targeting the indicated DUBs. Ub-MGFP and Ub-RGFP were transfected 48 h later and cells grown for 20 hrs post-transfection. Total cell lysates were prepared and analysed by Western blotting using GFP antibodies and tubulin (loading control). **B)** Ub-RGFP signals were quantified using the Odyssey ® Fc imaging system and normalised to the tubulin loading control, values show standard errors for n = 3. Pairwise comparisons relative to the NT control, were performed using GraphPad Prism. Student t-test: P > 0.05 = ns, P ≤ 0.05= *, P ≤ 0.01= **, P ≤ 0.001= ***. **C)** HeLa TRex Flp-In cells stably expressing OpD after induction were seeded at 50 % confluence and transfected with 1 nM of siRNAi targeting the indicated DUBs. OpD expression was induced 48 h later, and cells grown for 20 hrs post induction. Total cell lysates were prepared and analysed by Western blotting using opsin antibodies to detect OpD, and tubulin (loading control). **D)** OP91 signals were quantified using Odyssey ® Fc imaging system and normalised to the tubulin loading control, values show standard errors for n = 3. Pairwise comparisons relative to the NT control, were performed using GraphPad Prism. Student t-test: P > 0.05 = ns, P ≤ 0.05= *, P ≤ 0.01= **, P ≤ 0.001= *** and P ≤ 0.0001=****.

To further explore the substrate specificity of USP5, we looked at the effects of knocking down USP5 on the stability of the opsin degron (OpD), an ERAD substrate derived from wild-type opsin by the introduction of a degron motif into the first transmembrane domain [56]. The SGTA/ Bag6 complex was previously shown to play an important role in ERAD of OpD [20],, [47]. Knocking down USP5 did not change the steady state levels of OpD when compared to the non-targeting control (Fig. 3C, cf. lane 1 and 2). On the other hand, knocking down USP14 and USP9X resulted in an accumulation of OpD (Fig. 3C, cf. lane 1, 3 and 4). These effects mirrored what was observed when the proteasome is inhibited by Bortezomib (Fig. 3C, lane 5). This observation is consistent with a perturbation of ERAD upon knocking down USP14 or USP9X but not USP5. Based on these results, we concluded that USP5 is a DUB that specifically plays a role in the cytosolic quality control of MLPs.

### USP5 protects OP91 against proteasomal degradation

Having established that knocking down USP5 reduces the steady state levels of OP91. We next investigated whether knocking down USP5 changes the OP91 mRNA levels. HeLa T-REx cells were transfected with oligonucleotides to knockdown USP5 or a non-targeting control. After 72 hours, cells were harvested and total RNA was extracted, followed by analysis of OP91 mRNA levels using quantitative RT-PCR. In contrast to the observed decrease in steady state levels of the OP91 protein, mRNA levels were comparable between the non-targeting control and a USP5 knockdown (Fig. 4A), suggesting that the reduction in OP91 levels in a USP5 knockdown occurs post-translationally.

**Fig 4.**
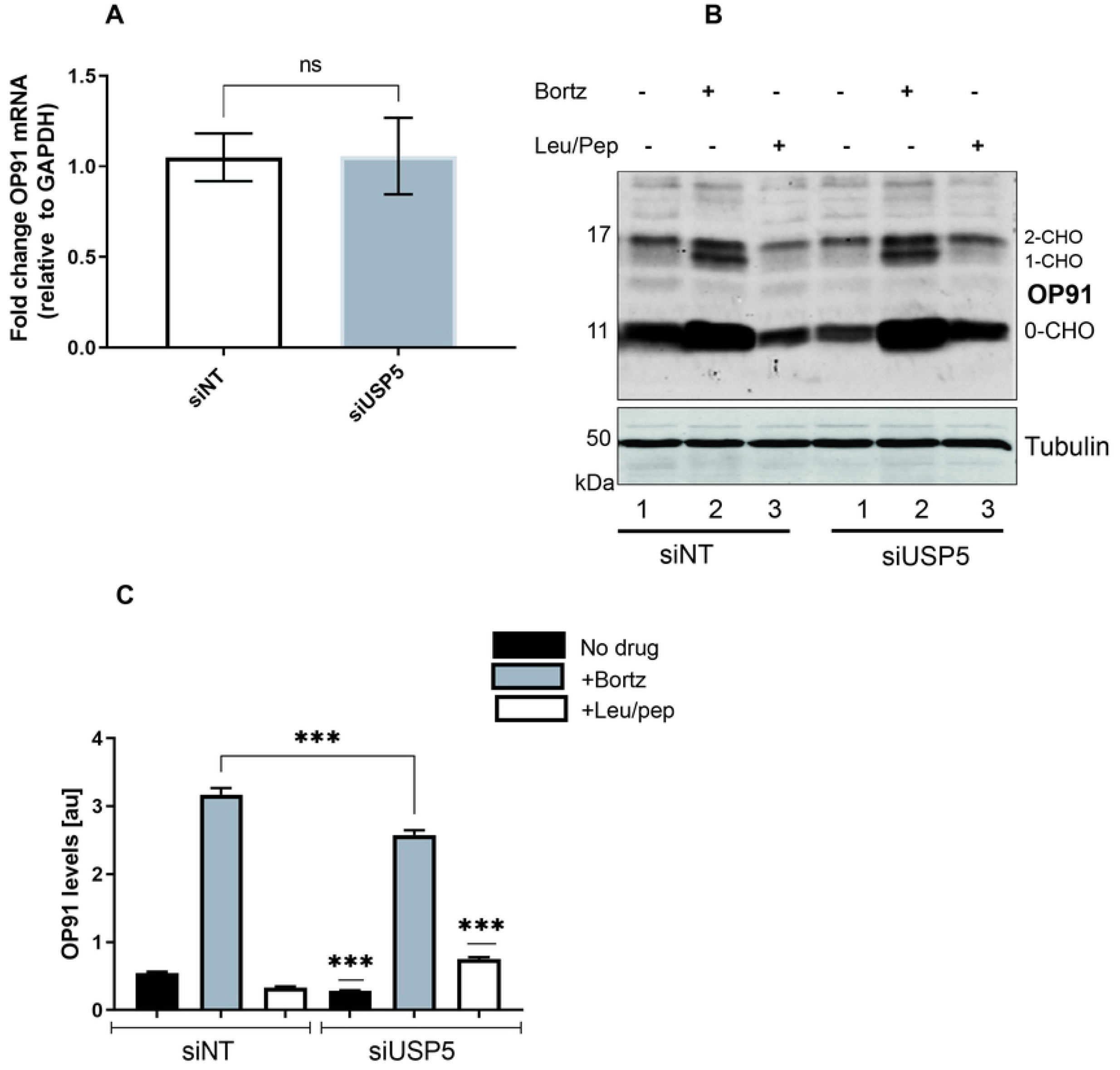
USP5 protects OP91 against proteasomal degradation. **A)** HeLa T-REx Flp-In cells stably expressing OP91 were seeded at 50% confluence followed by transfection of oligos for knockdown of USP5 or a non-targeting oligo. Cells were grown for 48 hrs followed by induction of OP91 expression and growth for a further 20 hrs post. Cells were harvested and total RNA extracted and used in a qPCR reaction following standard qPCR protocol. **B)** Cells grown as described in A, inhibitors of the proteasome and autophagy were added 24 hours before cells were harvested. Samples were analysed by SDS-PAGE and Western blotting for OP91 and Tubulin loading control. **C)** OP91 signals were quantified using Odyssey ® Fc Imaging System and normalised to the tubulin loading control, values show standard errors for n = 3. Pairwise comparisons were performed using GraphPad Prism. Student t-test: P > 0.05 = ns, P ≤ 0.05= *, P ≤ 0.01= **, P ≤ 0.001= *** and P ≤ 0.0001=****.

OP91 was previously shown to be degraded by the proteasome [32]. Therefore, we investigated whether OP91 was stabilised by proteasome inhibition when USP5 levels are reduced. Upon addition of Bortezomib, a proteasome inhibitor, there was an increase in steady state levels of OP91 in both the nontargeting control and in a USP5 knockdown (Fig. 4B cf. lanes 2 and 5). Although, Leupeptin/Pepstatin which inhibits autophagy did not increase the steady state levels of OP91 in the non-targeting control (Fig. 4B cf. lanes 3), this treatment led to stabilisation of a small proportion of OP91 in a USP5 knockdown (Fig. 5B cf. lane 6). This observation suggests that in the absence of USP5 a small pool of OP91 is degraded by autophagy, consistent with the cross talk observed between the proteasomal and autophagic pathways of degradation [57].

**Fig 5.**
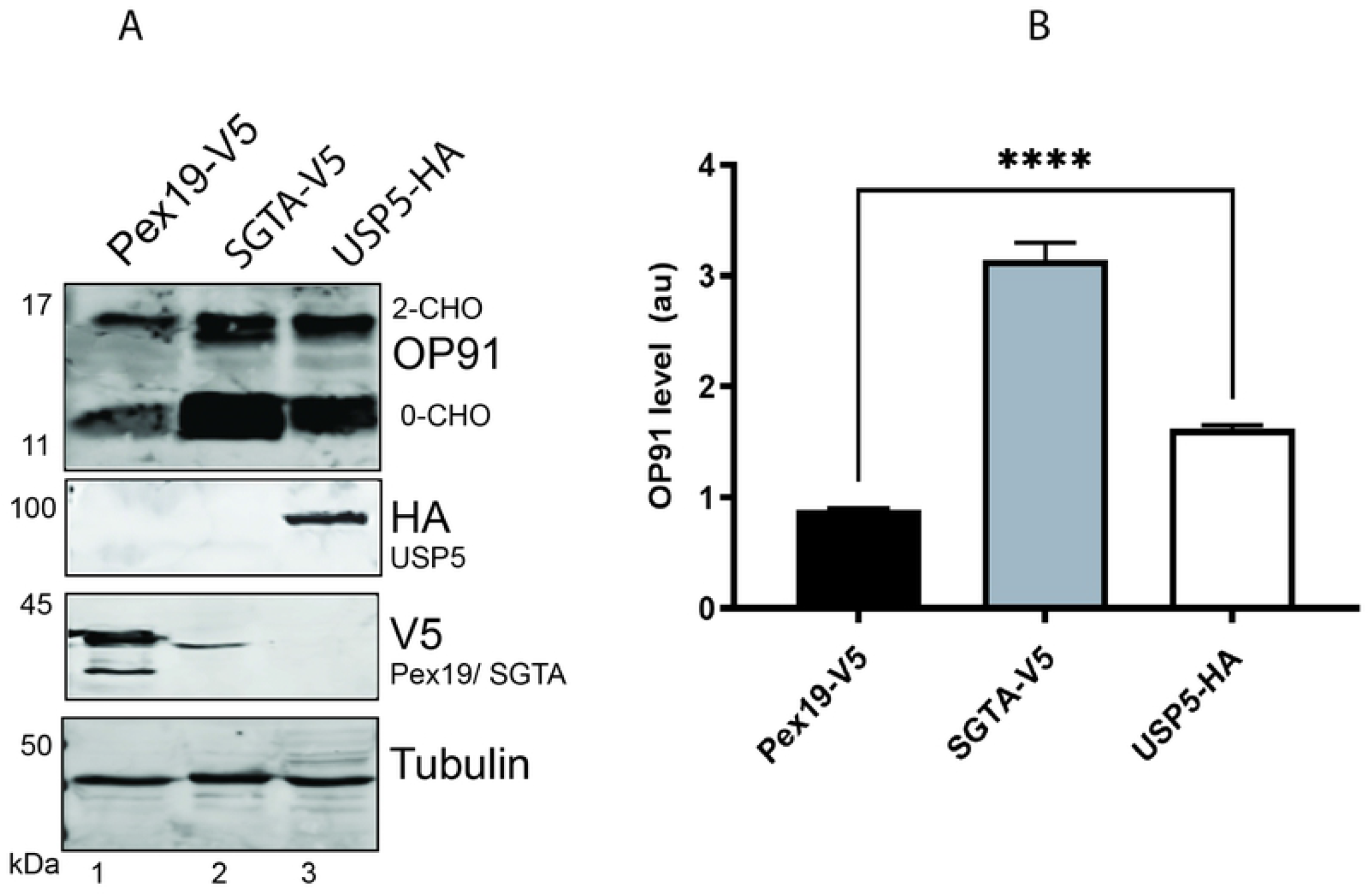
Overexpression of USP5 stabilises OP91. **A)** HeLa T-REx Flp-In cells stably expressing OP91 upon induction were transfected with V5-tagged Pex19 and SGTA or HA -tagged USP5. OP91 expression was induced 6 hours post-transfection. Cells were grown for a further 20 hours, post-induction. Total cell lysates were prepared and analysed by western blotting using antibodies against OP91, V5 tagged SGTA and Pex19, HA tagged USP5 and tubulin (loading control). **B)** OP91 signals were quantified using Odyssey ® Fc Imaging System and normalised to the tubulin loading control, values show standard errors for n = 3. Pairwise comparisons relative to Pex19, were performed using GraphPad Prism. Student t-test: P > 0.05 = ns, P ≤ 0.05= *, P ≤ 0.01= **, P ≤ 0.001= *** and P ≤ 0.0001=****.

To better understand the basis for the reduced steady-state levels of MLPs observed in a USP5 knockdown, we analysed OP91 levels over a 1-hour time course by blocking protein synthesis with cycloheximide and following degradation of OP91 in knockdowns of USP5, USP9X and in a nontargeting control. Previous studies showed that knocking down SGTA leads to a reduction in OP91 levels (S1. Fig cf. lane 2). We reasoned that if USP5 and SGTA act in the same pathway, their knockdown should affect the degradation of MLPs in a similar way. Therefore, an SGTA knockdown was included in the time-course experiments for comparison. Pulse chase experiments revealed that in the non-targeting and USP9X knockdowns, ∼50 % of OP91 was degraded in the first 20 minutes with the rate of degradation slowing after this point (see supplementary material, S2A. Fig). While, in the SGTA and USP5 knockdowns, there was a less marked reduction as the steady state levels of OP91 were already low before the start of the chase experiment (S2A. Fig). Therefore, to better compare the degradation kinetics in these knockdowns, OP91 levels were expressed as a percentage of the total OP91 signal in the negative control (NT) at time zero (S2B. Fig). This revealed similar degradation kinetics for all conditions, notably however, these experiments also revealed that when USP5 and SGTA are reduced there is loss of a protected pool of OP91 which is rapidly degraded.

### Overexpression of USP5 stabilises OP91

To gain confidence of the role of USP5 in MLP quality control, we investigated the effect of overexpressing USP5 on the stability of OP91. To this end HA tagged USP5, and V5 tagged SGTA [43] were transfected into HeLa cells followed by induction of OP91 expression. We have previously shown that overexpression of a chaperone protein, Pex19 does not stabilise OP91 [43]. Therefore, this was used as a negative control to reveal the basal level of OP91. After growth, cells were harvested into Laemmli sample buffer and analysed by Western blotting, probing for OP91, Tubulin, USP5, SGTA and PEX19. Consistent with previous studies (Wunderley, Leznicki et al. 2014), overexpression of SGTA led to an increase in the steady state levels of OP91 (Fig 5A cf. lane 2), above those seen in the presence of exogenous PEX19 control (Fig. 5A cf. lane 1). Overexpression of USP5 led to 2-fold increase in the steady state levels of OP91 when compared to the Pex19 control (Fig. 5A cf. lane 3). These results demonstrated that modulating USP5 levels directly affect MLP quality control.

### USP5 specifically enhances SGTA activity

It was previously shown that increasing exogenous SGTA levels leads to an increase in steady state levels of OP91 [32],[43]. Having shown that USP5 is also required for the stabilisation of OP91 in this study (Fig. 2A & B), we next asked whether the stabilising effect of SGTA is dependent on USP5. To this end we performed a knockdown of USP5 and used non-targeting siRNAi as control. After 48 hours, V5 tagged SGTA or Pex19 control were transfected followed by induction of OP91 expression. After growth, cells were analysed by Western blotting for OP91, tubulin, and V5 tagged SGTA or Pex19. Overexpression of SGTA led to stabilisation of OP91 in the presence of the nontargeting siRNAi (Fig. 6A cf. lane 2), when compared to the Pex19 control (Fig. 6A cf. lane 1). However, in the presence of USP5 siRNAi, overexpression of SGTA did not stabilise OP91 to the same level as observed in the presence of non-targeting siRNAi (Fig. 6A cf. lane 2 and 4), suggesting that USP5 is required for the stabilisation of OP91 observed during SGTA overexpression.

**Fig 6.**
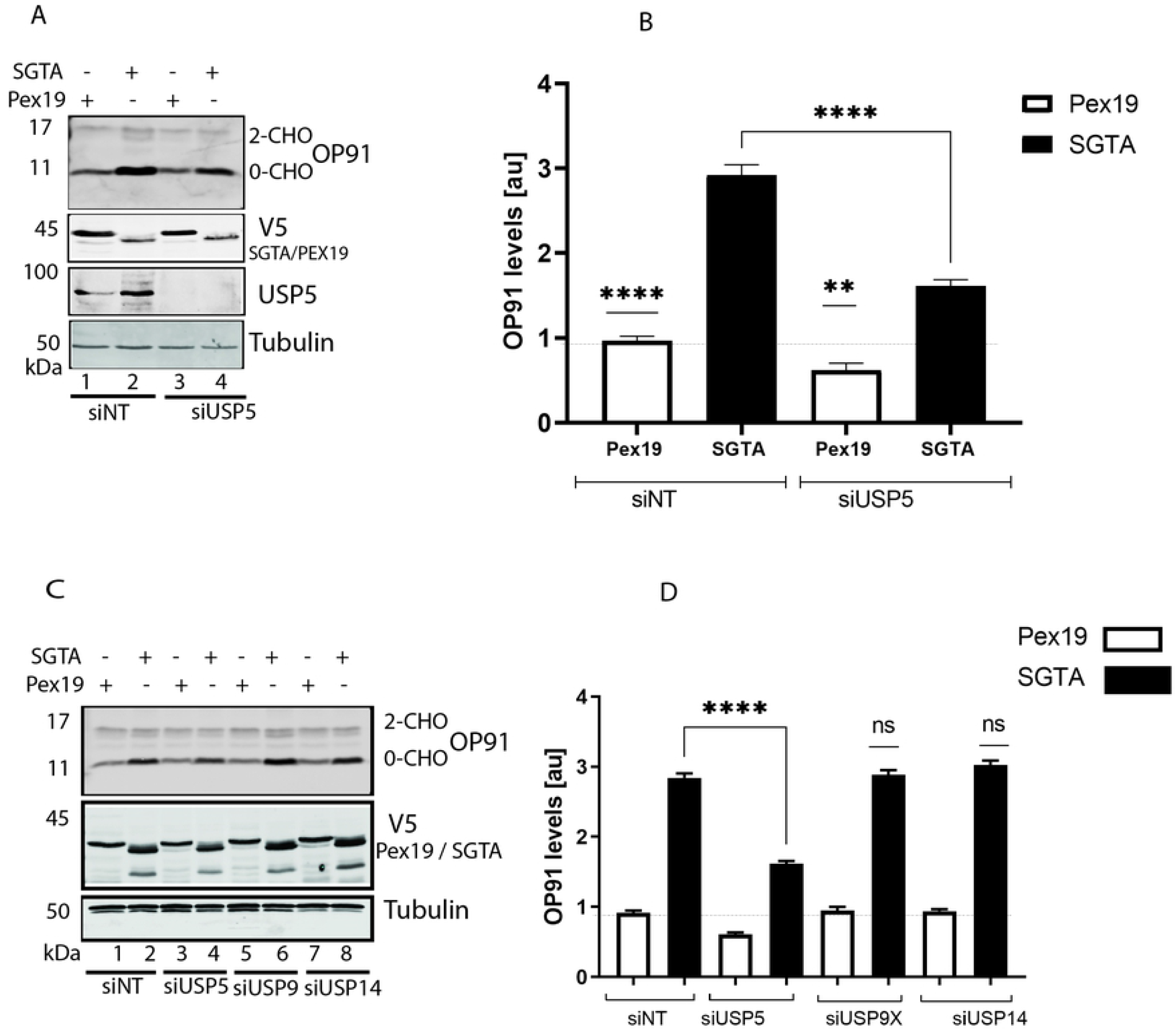
USP5 specifically enhances SGTA activity. **A)** HeLa T-REx Flp-In cells stably expressing OP91 after induction were seeded at 50 % confluence and transfected with 1 nM of siRNAi targeting USP5 or a nontargeting control and incubated for 48 hours. After this, plasmids expressing V5 tagged SGTA or PEX19 were transfected followed by incubation for 6 hours before the medium was replaced with medium containing 1 mg/ml of tetracycline to induce OP91 expression. Cells were grown for 20 hrs post induction and total cell lysate were prepared and products analysed by western blotting using antibodies for opsin (OP91), V5 (SGTA and PEX19), and tubulin (loading control). **B)** OP91 signals were quantified using Odyssey ® Fc Imaging System and normalised to the tubulin loading control, values show standard errors for n = 3. Pairwise comparisons relative to Pex19, were performed using GraphPad Prism. Student t-test: P > 0.05 = ns, P ≤ 0.05= *, P ≤ 0.01= **, P ≤ 0.001= *** and P ≤ 0.0001=****. **C)** HeLa T-REx Flp-In cells were treated with siRNAi targeting USP5, USP9X, USP14 and a nontargeting control and analysed as detailed in A. **D)** OP91 signals were quantified using Odyssey ® Fc Imaging System and normalised to the tubulin loading control, values show standard errors for n = 3. Pairwise comparisons relative to Pex19, were performed using GraphPad Prism. Student t-test: P > 0.05 = ns, P ≤ 0.05= *, P ≤ 0.01= **, P ≤ 0.001= *** and P ≤ 0.0001=****.

We also revisited the effect we observed that upon knockdown of USP14 destabilisation of OP91 occurred (Fig. 2A). To investigate whether USP14 is required for the stabilisation effect observed with exogenous SGTA, we knocked down USP14 and overexpressed SGTA. We also looked at the effects in a USP9X knockdown as this did not affect the steady state levels of OP91 (Fig 2A & B). As expected, reducing USP9X did not affect the ability of SGTA to stabilise OP91 (Fig. 6C cf. lane 5 and 6). On the other hand, lack of USP5 led to a reduction in the ability of SGTA to stabilise OP91 (Fig. 6C cf. lane 3 and 4). However, the knockdown of USP14 did not affect the ability of exogenous SGTA to stabilise OP91 (Fig. 6C cf. lane 7 and 8), suggesting that USP14 is not directly required for the SGTA mediated stabilisation of OP91.

### Association of SGTA and USP5 is enhanced in the presence of an MLP

Co-immunoprecipitation studies have shown that SGTA and OP91 are in a complex [32]. We next wanted to see whether USP5 was also part of this complex. To this effect, HeLa cells were transfected with SGTA, followed by induction of OP91 expression in half of the samples. After growth, cells were harvested into non-denaturing buffer and subjected to co-immunoprecipitation using SGTA antibodies or IgG antibodies as negative control. SGTA associated with USP5 in the absence of OP91 induction and this association was increased upon OP91 induction (Fig. 7A cf. lane 5 and 6), suggesting that association of USP5 and SGTA is substrate dependent. This association was more pronounced for the non-glycosylated form [0-CHO] of OP91 which probably reflects the mislocalised form. Since the IgG control did not show an increased association of USP5 with SGTA even upon OP91 induction (Fig. 7A cf. lane 3 and 4), these results suggested that this association was specific for SGTA.

**Fig 7.**
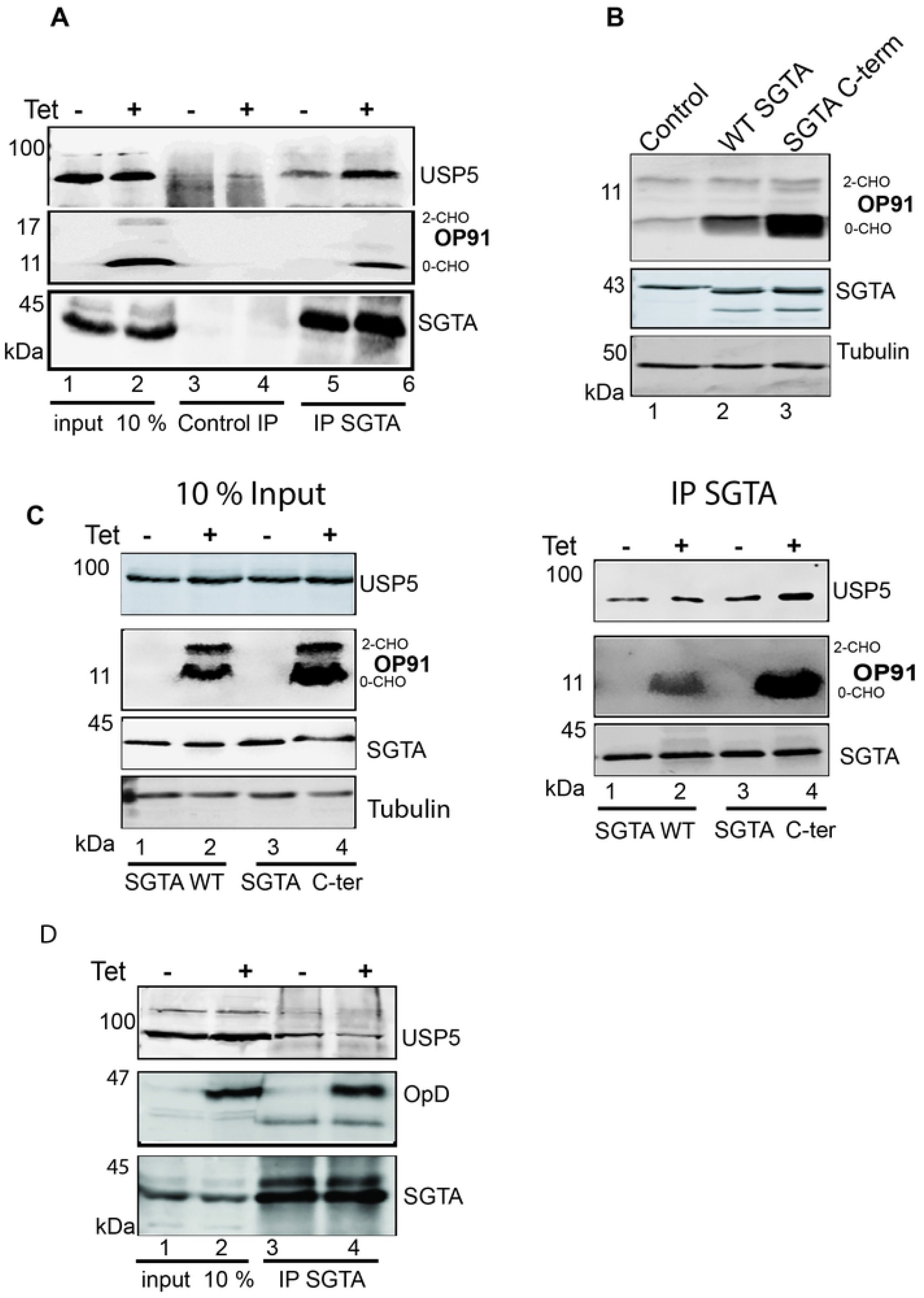
Association of SGTA and USP5 is enhanced in the presence of an MLP. **A)**HeLa T-REx Flp-In cells stably expressing OP91 after induction were seeded at 50 % confluence and transfected with SGTA, followed by induction of OP91 expression in half of the samples. After growth, cells were harvested into non-denaturing buffer and subjected to co-immunoprecipitation using SGTA antibodies or IgG antibodies as negative control. Products were analysed by western blotting using antibodies against opsin (OP91), V5 (SGTA) and tubulin (loading control). **B)** HeLa TRex Flp-In cells stably expressing OP91 after induction were transfected with V5 tagged WT SGTA or SGTA-3xNNP/AAA C-terminal mutant (denoted here as SGTA C-term) followed by incubation for 6 hours before the medium was replaced with medium containing 1 mg/ml of tetracycline to induce OP91 expression. Cells were grown for 20 hrs post induction and total cell lysate were prepared and products analysed by western blotting using antibodies for opsin (OP91), V5 (SGTA and SGTA-C-term), and tubulin (loading control). **C)** Cells transfected with WT SGTA or the SGTA C-terminal mutant described in B were processed for immunoprecipitation using SGTA antibodies. Samples were analysed by western blotting using antibodies against SGTA, OP91 and USP5. **D)** HeLa TRex Flp-In cells stably expressing OpD upon induction were seeded at 50 % confluence and transfected with SGTA, followed by induction of OpD expression in half of the samples. Cells were subjected to immunoprecipitation as detailed in A.

SGTA contains a well conserved C-terminal NNP region characterized by three repeats of the NNP motif. Previously, a mutant version of SGTA in which all three NNP motifs were altered to alanine’s (SGTA-3xNNP/AAA-V5), was shown to enhance the ability of SGTA to stabilise OP91 five -fold [46], (Fig 7B, cf. lane 2 and 3). To further investigate the substrate dependent association of SGTA and USP5 we made use of this SGTA mutant. We reasoned that if the stabilisation of MLPs is linked to deubiquitination, this mutant will likely associate with more USP5 than WT SGTA. We therefore asked whether this mutant associates with more USP5 compared to the WT SGTA. To this end, HeLa cells were transfected with WT SGTA or the SGTA mutant, followed by induction of OP91 expression in half of the samples. Cells were harvested into non-denaturing buffer 24 hours post-induction and subjected to co-immunoprecipitation using SGTA antibodies or IgG antibodies as a negative control. As expected SGTA associated with USP5 in the absence of OP91 induction and this association was increased upon OP91 induction (Fig. 7C cf. lane 3 and 4). Similarly, the SGTA-3xNNP/AAA mutant associated with more USP5 in the absence of induction (Fig. 7C cf. compare lane 1 and 3), consistent with an increase in stabilisation of OP91 seen in this mutant before induction. Upon induction of OP91, there was an increase in association compared to the WT (Fig. 7C cf. compare lane 2 and 4), concomitant with an increase in OP91. Therefore, these results confirmed that this association is substrate dependent.

To validate these observations, we investigated the association of SGTA and USP5 in the presence of the ERAD substrate, opsin degron (OpD) [20], [47]. Induction of OpD did not result in an increase in the association of SGTA and USP5 when compared to the uninduced sample (Fig. 7D, cf. lane 3 and 4). Based on these results, we concluded that association of USP5 and SGTA is specifically enhanced by the presence of an MLP. Attempts to confirm direct binding of SGTA to USP5 *in vitro* did not show direct binding, suggesting the interaction may be via an adaptor protein.

## Discussion

A role for DUBs in SGTA mediated protein quality control was first apparent when SGTA was reported to promote deubiquitination of MLPs [9, 32]. However, until now the deubiquitinating enzymes that play a role in this process had not been identified. In this study, we identified DUBs associated with SGTA by using pull-down assays and mass spectrometry. We confirmed that knocking down USP5 specifically reduces the steady state levels of a model MLP (Fig. 2), while overexpression of USP5 leads to an increase (Fig. 5). Using co-immunoprecipitation, we show an association of USP5 and SGTA which is dependent on the presence of an MLP: association of USP5 with SGTA was increased upon induction of OP91 (Fig. 7). This association is specific for MLPs as we did not observe an increase in the association of USP5 with OpD, an ERAD substrate (Fig. 7). Moreover, a gain of function mutant of SGTA which leads to a 20-fold increase in the steady state levels of OP91 (Fig.7) showed increased USP5 association when compared to WT SGTA (Fig. 7). The failure of SGTA to effectively stabilise OP91 in the absence of USP5 (Fig. 6) points to an important role of USP5 in SGTA mediated quality control of membrane proteins that mislocalise to the cytosol. This is supported by an increased association of USP5 with the non-glycosylated OP91 which is mislocalised but has not been ER translocated (Fig.7). Overall, our results point to a role of USP5 in regulating the quality control of MLPs.

USP5 has a well characterised function regulating a stable pool of free ubiquitin in vivo by recycling unanchored polyubiquitin chains [13]. In this regard, USP5 hydrolyses K63-, K48-, K11-, K29-linked and linear polyubiquitin chains [58]. However, a role for USP5 in the stabilisation of substrates destined for degradation is not without precedence. Previously, overexpression of USP5 was shown to stabilise FoxM1, a protein with a key role in tumorigenesis and progression of pancreatic cancers [59]. In addition, USP5 was shown to associate with the linker III-IV of the Cav3.2 protein and its knockdown increased ubiquitination of Cav3.2 protein and subsequent degradation, thus supporting the notion that USP5 stabilises Cav3.2 [60].

USP5 shares 54.8% sequence identity with USP13, hence a partial functional overlap between the two has been suggested. Indeed, USP5 and USP13 are both recruited to stress granules containing K48 and K63 linked ubiquitin chains. In this case, depletion of USP5 and USP13 led to an increase in ubiquitinated proteins in heat induced stress granules, suggesting these DUBs play a role in ubiquitin editing of substrates to facilitate disassembly of stress granules [61]. Despite these functional overlaps, there is evidence supporting the idea that these closely related DUBs also have some distinct roles and classes of substrates. For instance, USP13 was previously shown to promote a stable interaction between BAG6 and SGTA during ERAD and its knockdown reduced this interaction [20]. On the other hand, USP5 did not have a direct effect on the interaction of SGTA and BAG6 [20]. Consistent with these observations, in this study we did not observe an effect of USP5 on the stability of the opsin degron, an ERAD substrate. Moreover, in our pulldown assays, we did not detect an association of SGTA with USP13, suggesting USP13 might have a limited or no role in MLP quality control.

Our results clearly indicate that USP5 enhances SGTA mediated MLP quality control: the ability of SGTA to stabilise MLP was significantly reduced in the absence of USP5 (Fig. 6). However, it is not clear how USP5 exerts these effects. This is complicated by the fact that DUBs such as USP5 have many cellular targets. Therefore, the effects of USP5 may be indirect as was reported for USP13 [21]. It was shown that, USP13 prevents promiscuous gp78-mediated ubiquitination of Ubl4A, a BAG6 cofactor which promotes ERAD. Ubiquitination of Ubl4A was accompanied by irreversible cleavage of BAG6 and attenuation of ERAD [21]. Hence, the deubiquitination action of USP13 eliminates ubiquitin conjugates from Ubl4A to maintain the function of BAB6, thereby ensuring efficient disposal of ERAD substrates. Recently, it was also shown that USP13 and gp78 also regulate the caspase activity of CASP3, which cleaves BAG6 and switches its function from an ERAD regulator to an autophagy tuner [19]. It was shown that the LIR1 motif of BAG6 holds the autophagosome structural membrane proteins LC3B-I and the unprocessed form of LC3B to suppress autophagy. This inhibition is relieved by the ubiquitination of thioredoxin which results in CASP3 activation and cleavage of BAG6 [19]. Hence, it may be that USP5 might exerts its’ effects by modulating the ubiquitination status of a key player in the degradation pathway of MLPs rather than the MLP itself. Reconstitution of the DUB reaction *in vitro* is necessary in order to fully defined the role of USP5 in MLP quality control.

Functional pairing of E3 ligases with DUBs, as discussed for gp78 and USP13, is an important regulatory step in protein quality control [15]. In MLP quality control, BAG6 binds hydrophobic substrates and recruits RNF126, an E3 ubiquitin ligase which catalyses the selective ubiquitination of MLPs [31, 44]. On the other hand, DUBs such as USP5 promote substrate deubiquitination thereby giving substrates a chance to avoid proteasomal degradation [9, 32, 43]. Hence, future work should study the interplay of USP5 and RNF126 in regulating MLP quality control. Another possibility is that USP5 might play a key role in ubiquitin chain editing, by promoting K-63 linked assembly of substrates which would favour their accumulation in inclusion bodies. This would be the reverse of what is seen with the DUBs, Ubp2 and Ubp3 which prevent assemble of K-63 chains on substrates thereby promoting K-48 mediated proteasomal degradation [18]. To understand the role of USP5 in MLP quality control, it is necessary to study the type of ubiquitin linkages of MLPs and how these change upon rapid depletion of USP5 by using approaches such as the auxin-inducible degron (AID) technology [62].

In this study we did not detect a direct interaction of USP5 and SGTA, suggesting the interaction might be via an adaptor protein. A key role of the adaptor protein UBXN1 in regulating the interaction of VCP with BAG6 clients occurring prior to ER insertion was recently demonstrated [63]. It was shown that BAG6 clients, once ubiquitylated, were not directly degraded by the proteasome, but were recognised by the VCP-UBXN1 complex prior to degradation. Failure of ubiquitinated substrates to engage the UBXN1 adaptor led to their accumulation as aggregates [63], underscoring the importance of the adaptor proteins in protein quality control. In the case of USP5, the dendritic cell-derived ubiquitin (Ub)-like protein (DC-UbP) functions as an adaptor protein reconciling the cellular ubiquitination and deubiquitination processes by regulating the function of USP5 and UbE1 [64]. The C-terminal UbL domain of DC-UbP interacts with the UBA domains of USP5 and with the Ub-fold domain (UFD) of the Ub-activating enzyme UbE1, on the distinct surfaces. Overexpression of DC-UbP enhanced the association of these two enzymes and prompted cellular ubiquitination, whereas knockdown of the protein reduced the cellular ubiquitination level [64]. In the case of MLP quality control, SGTA may function in a similar way to DC-UbP as it has a ubiquitin (Ub)-like domain on the N-terminus where BAG6 binds and promote RNF126 recruitment [31]. It may be that this (Ub)-like domain also promotes recruitment of DUBs such as USP5, via another protein. Future work should investigate the nature of interaction between USP5 and SGTA and identify any adaptor proteins involved in this process.

The number of DUBs that play important roles in the cytosolic protein quality control continues to increase, and this study adds USP5 to this list. This study suggests that USP5 impacts on the disposal of MLPs and possibly other aggregation prone proteins when its levels are modulated. Therefore, future work should extend these studies to other membrane proteins and to substrates implicated in pathologies such as neurodegeneration and cancer in order to understand how USP5 regulates their fates. These studies will shed light on how USP5 can be modulated as a therapeutic target for prevention of terminal aggregates which are linked to neurodegenerative diseases, cancer, and type II diabetes II. The apparent specificity of USP5 for cytosolic substrates, makes effective therapies more likely as there may be a limited crosstalk to other DUBs and pathways.

## Acknowledgements

We thank, Prof. Stephen High for his guidance and support with reagents and manuscript preparation, without his input this work would not have been possible. Dr Pawel Leznicki for proofreading and advice during the preparation of this manuscript.

## Supporting information

**Fig_S1. Knockdown of SGTA destabilises OP91.**

HeLa T-REx Flp-In cells stably expressing OP91 after induction were seeded at 50 % confluence and transfected with 1 nM of siRNAi targeting SGTA or a nontargeting control and incubated for 48 hours. After this, OP91 expression was induced by replacing with medium containing 1 mg/ml of tetracycline. Cells were grown for 20 hrs post induction and total cell lysate were prepared and analysed by western blotting using antibodies for opsin (OP91), SGTA, and tubulin (loading control) and detected using an Odyssey ® Fc imaging system.

**Fig_S2. USP5 and SGTA protect OP91 from degradation.**

**A.** HeLa T-REx Flp-In cells stably expressing OP91 were seeded at 50 % confluence and transfected with 1 nM of siRNAi targeting USP5, USP9X, SGTA or a nontargeting control and incubated for 48 hours. After this OP91 expression was induced by addition of medium containing 1 µg/ml of tetracycline and grown for 20 hrs post induction. Prior to harvesting transfected cells were treated with 100 µg/ml cycloheximide (Sigma, Aldrich) to inhibit protein synthesis. Cells were lysed directly into SDS-PAGE sample buffer at specific time-points followed by Western blotting with antibodies against opsin (OP91) and tubulin (loading control) using fluorescence-based detection (LICOR). **B)** OP91 signals were quantified using Odyssey ® Fc imaging system and normalised to the tubulin loading control, values show standard errors for n = 3.

